# Brain-wide mapping of neuronal architecture controlling torpor

**DOI:** 10.1101/2023.03.03.531064

**Authors:** Hiroshi Yamaguchi, Keith R Murphy, Noriaki Fukatsu, Kazuhide Sato, Akihiro Yamanaka, Luis de Lecea

## Abstract

Endotherms can survive low temperatures and food shortage by actively entering a hypometabolic state known as torpor. Although the decrease in metabolic rate and body temperature during torpor is controlled by the brain, the underlying neural circuits remain largely unknown. In this study, we identify the neural circuits involved in torpor regulation by combining whole-brain mapping of torpor-activated neurons, cell type-specific manipulation of neural activity, and viral tracing-based circuit mapping. We found that Trpm2-positive neurons in the preoptic area and Vgat-positive neurons in the dorsal medial hypothalamus are activated during torpor. Genetic silencing shows the activity of either cell type is necessary to enter the torpor state. Finally, we show that these cells receive projections from the arcuate and suprachiasmatic nucleus and send projections to brain regions involved in thermoregulation. Our results demonstrate an essential role of hypothalamic neurons in the regulation of body temperature and metabolic rate during torpor and identify critical nodes of the torpor-regulatory network.

## Introduction

Endotherms can adapt to a wide range of environmental temperatures by producing metabolic heat and maintaining a constant body temperature, but require more food as a source of energy. Some endotherm enter a hypoactive metabolic state known as torpor when food shortage or low environmental temperature make it difficult to maintain body temperature. During torpor, animals suppress metabolic rate by 70 - 90% and lower body temperature by 10 - 30°C. Across over 200 species, including primates, torpor can be classified as seasonal torpor (hibernation)^12^, which lasts from days to several weeks or short torpor (daily torpor) which cycles within a day ^23^. Previous works found that displacement of hypothalamic temperature during torpor alters body heat production^4^, and that central administration of adenosine receptor antagonists blocks entry into torpor in hibernating animals^5,6^ and mice^7^. While these findings suggest the brain controls torpor, the exact circuits involved are largely unknown. Here we induced torpor in laboratory mice by fasting and performed brain-wide mapping of torpor-activated neurons, cell-type specific manipulation of neural activity, and viral tracing to interrogate the neuronal circuits that control torpor.

## Results

### Torpor induction in mice

We first measured the body temperature (Tb) and carbon dioxide production (VCO_2_) of mice for 24 hours under four conditions: fed at thermoneutrality (31°C), fasted at thermoneutrality, fed at 16°C or fasted at 16°C. The experiment was started at the beginning of the dark phase in a 12-12 hours light/dark cycle. We found that, rather than chronically entering hypotheremic and metabolic state, the shift exhibited by mice fasted at 16 °C was rapid and reversible and could occur several times within a day(Fig. 1A). To better quantify the torpor state, we examined the maximum decrease in Tb relative to a baseline period (ZT12-14, Δ) and found that mice fasted at 16 °C decrease Tb by 11.9 ± 0.7 °C, whereas other conditions had fairly moderate decrease of 2.4-3.1 °C (Fig. 1B). The VCO_2_ of cold-fasted mice decreased by 90 ± 1.4%(Fig. 1C), and preceded the decrease in Tb (Fig. 1A). In contrast, the maximum decrease under other conditions was 52 ∼ 72% (Fig. 1C). Torpid mice also dramatically decreased locomotor activity (figs. S1A) and took a rounded posture to minimize heat loss from the body surface (figs. S1B).

**Fig. 1.**
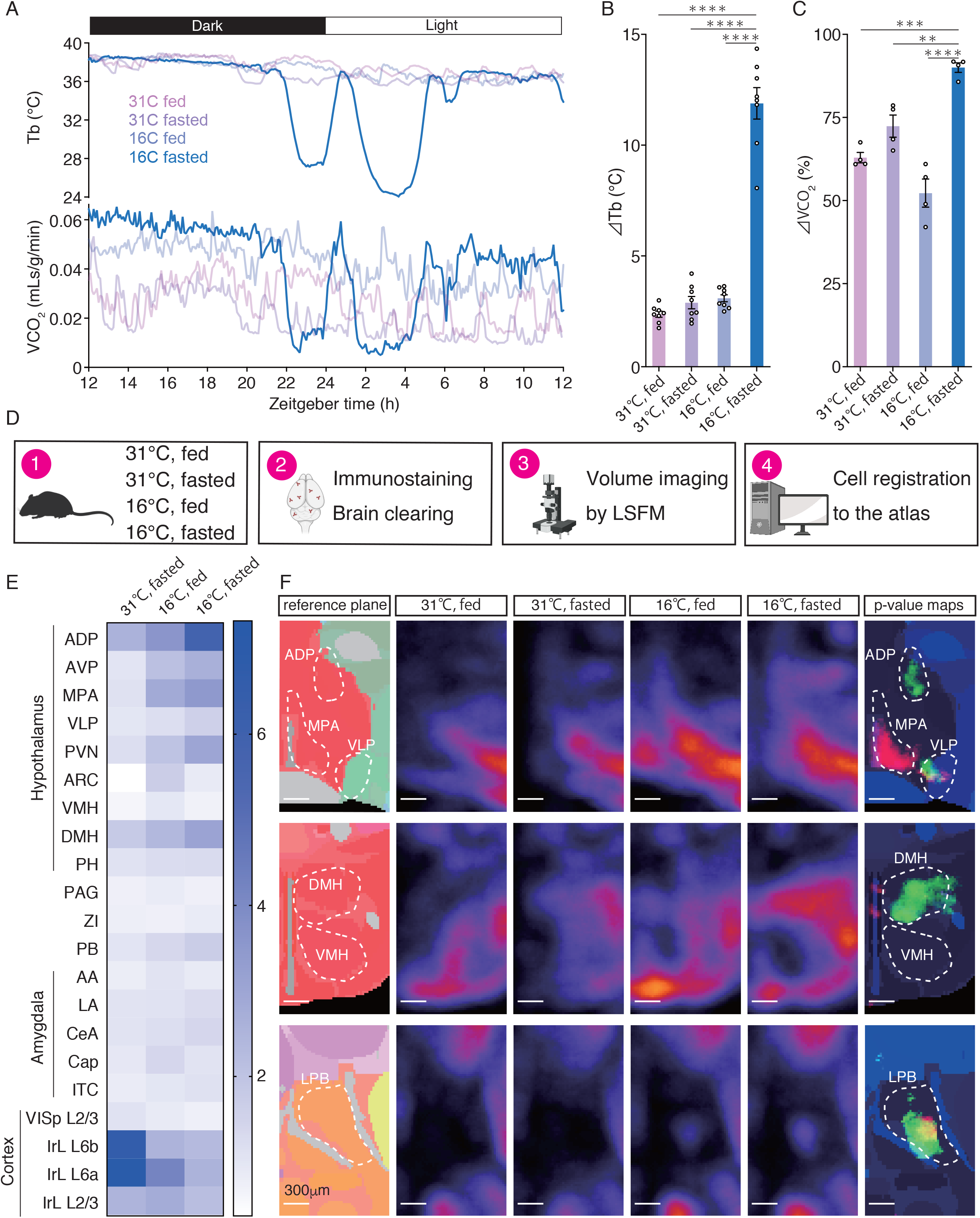
Brain-wide mapping of torpor-activated neurons. **(A)** Representative tracings of body temperatures (Tb) and CO_2_ productions (VCO_2_) in mice at thermoneutrality (31°C) or cold (16°C) with fed or fasted. **(B)** Quantification of the change in Tb at thermoneutrality (31°C) or cold (16°C) with fed or fasted (n=8 each). ⊿Tb was defined as the mean Tb for the first 120min minus the minimum Tb. **(C)** Quantification of the change in VCO_2_ at thermoneutrality (31°C) or cold (16°C) with fed or fasted (n=4 each). ⊿VCO_2_ was defined as the percentage reduction in VCO_2_ by comparing the mean VCO_2_ for the first 120min to the minimum VCO_2_. **(D)** Experimental design of brain-wide screen of torpor-activated neurons. Brain samples were collected from the mice at thermoneutrality (31°C) or cold (16°C) with fed or fasted (n=4 each) at ZT20. The brains were stained with anti-phospho-S6 antibody and optically cleared. The cleared brains were scanned by light sheet fluorescence microscopy (LSFM) and pS6+ neurons were mapped to Allen Brain Atlas. **(E)** Representative results of our brain-wide activity mapping. The groups under different conditions are normalized to the data of control mice fed at thermoneutrality. Color-coded scale bar indicates fold increases in the number of pS6+ neurons relative to control group for the following regions; Anterodorsal preoptic area (ADP), Anteroventral preoptic area (AVP), Medial preoptic area (MPA), Ventrolateral preoptic area (VLP), Paraventricular hypothalamic nucleus (PVN), Arcuate hypothalamic nucleus (ARC), Ventromedial hypothalamic nucleus (VMH), Dorsomedial hypothalamic nucleus (DMH), Posterior hypothalamic nucleus (PH), Periaqueductal gray (PAG), Zona incerta (ZI), Parabrachial nucleus (PB), Anterior amygdalar area (AA), Lateral amygdalar nucleus (LA), Central amygdalar nucleus (CeA), amygdalar capsule (Cap), Intercalated amygdalar nucleus (ITC), Primary visual area, layer 2/3 (VISp L2/3), Infralimbic area, layer 6b (IrL L6b), Infralimbic area, layer 6a (IrL L6a), Infralimbic area, and layer 2/3 (IrL L2/3). **(F)** Automated analysis of pS6+ cell distribution in mouse brains collected under different conditions at ZT20 (n=4). The number of pS6 + cells was counted per each voxel. Panels show the reference annotation, the averaged cell density maps for each group, and p-value maps for the following regions; ADP (top), MPA (top), VLP (top), DMH (middle), VMH (middle), and lateral parabrachial nucleus (LPB) (bottom). The cell density from each group were compared with that of control mice fed at thermoneutrality by the *t*-test and their p-values (Magenta: fasted at 31°C, Red: fed at 16°C, Green: fasted at 16°C) were mapped. Scale bar, 300μm.

### Whole-brain mapping of torpor-activated neurons

To identify the neuronal populations that control torpor, we performed a whole-brain screen for torpor-activated neurons with optically clear, intact brain volume imaging. Because mice fasted at cold ambient temperature usually begin torpor at ZT 19 (437 ± 28 min after onset of dark phase) (fig. S2), brains were harvested from mice at ZT 20 under four conditions (n = 4), permeabilized, immunolabeled with antibodies against the active neuron marker phospho-S6^8^, and cleared by the iDISCO method^9^. Immunoreactive cells in one hemisphere of the optically cleared brain were imaged using light sheet fluorescence microscopy and the cells were mapped to Allen brain Atlas^10^ using the ClearMap pipeline^11^ (Fig. 1D). Immunoreactive cells were counted in 1198 brain regions (Table 1) and normalized to the number of 31°C-fed controls to generate a brain activity map (Fig. 1E). We found that the antero-preoptic area (ADP and AVP), medial preoptic area (MPOA), paraventricular nucleus (PVN), and dorsomedial hypothalamus (DMH) were all strongly activated compared with other control brains (Fig. 1E). These hypothalamic regions have been reported to be involved in thermoregulation^12^, supporting the accuracy of our screening results. In contrast, neural activity decreased in torpor mice in various regions of the cerebral cortex (Fig. 1E). We further examined the number of immunoreactive cells in each single voxel of the reconstructed brain, and then generated p-value maps by comparing four groups of mice (Supplemental Movie. 1). The ventral part of the medial preoptic area was activated in mice fed at cold ambient temperature, but not in torpid mice, suggesting that this region is involved in thermogenesis and be suppressed during torpor (Fig. 1F top). The antero-preoptic area and ventrolateral preoptic area were specifically activated in torpid mice (Fig. 1F top). The anterior part of DMH was activated only in torpid mice (Fig. 1F middle). Activation of PB neurons in both groups fed or fasted at cold ambient temperature (Fig. 1F bottom) was consistent with a previous report that this region responded to lower ambient temperature^13^. In addition, the BNST and lateral septum in the torpid mice had more immunoreactive cells compared to the region in the control mice (fig. S3).

### Vgat+ Dorsomedial Hypothalamic neurons are active during and necessary for torpor

Among the regions associated with torpor, we focused on the DMH, which was among the most activated brain regions in torpid mice (Fig. 1E and 1F). A time course study identified a cluster of phospho-S6-positive neurons in the anterior DMH of torpid mice at ZT 20 which were largely absent in control mice (Fig. 2A). We also found that the DMH began to be activated at ZT18 before torpor onset. To characterize torpor-activated neurons in the DMH, we injected adeno-associated virus (AAV) carrying Cre-dependent eYFP into the DMH of Vgat-cre animals (Fig. 2B Left). Three weeks after virus injection, mice were fasted at 16 ° C and brains were collected 60 minutes after the onset of torpor. In the DMH from Vgat-cre mice, 75% of phospho-S6-positive cells were eYFP-positive (Fig. 2B right and 2C). Taken together, these results indicate that torpor-activated neurons in the DMH are predominantly GABAergic.

**Fig. 2.**
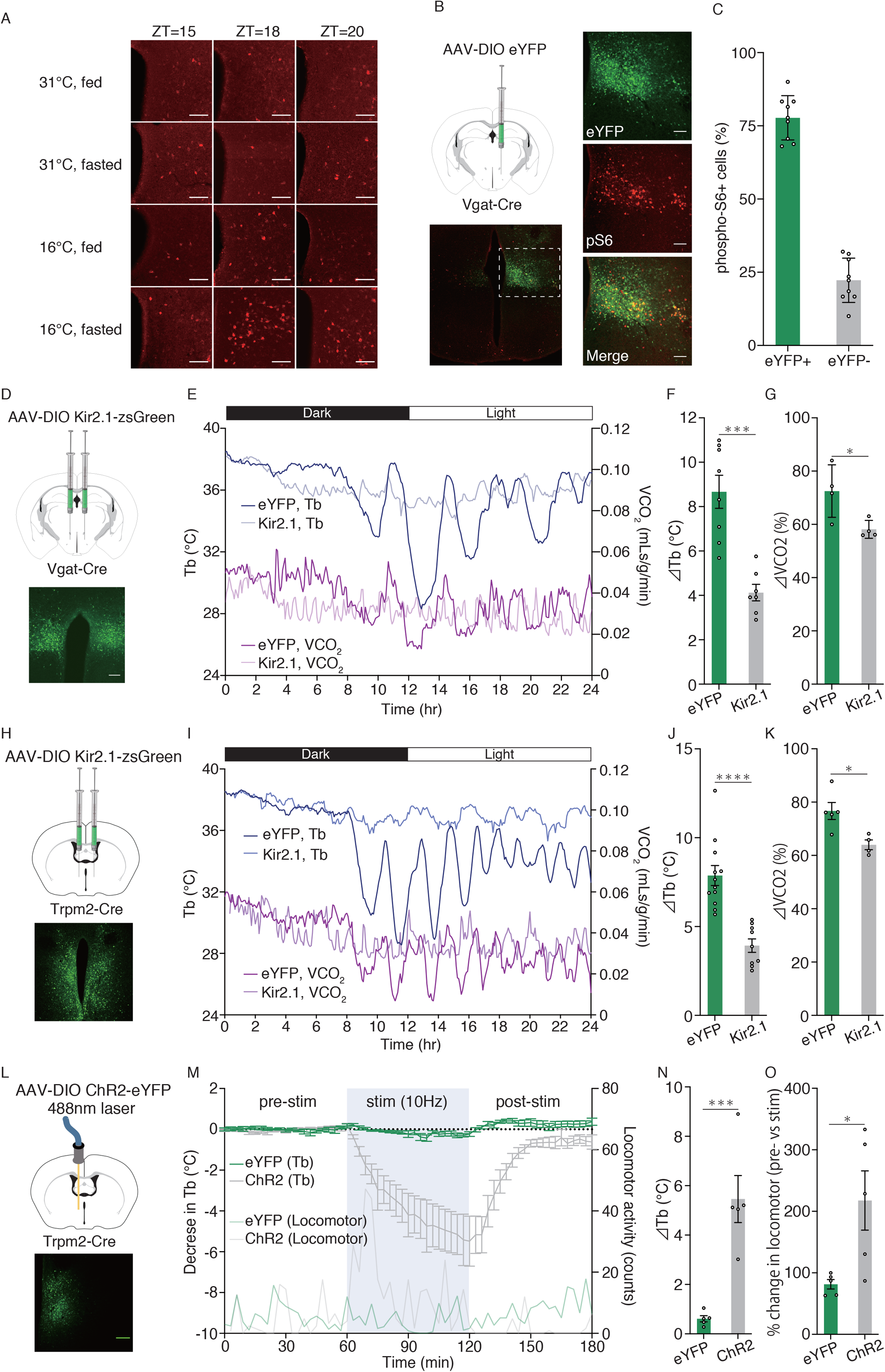
Hypothalamic neurons regulate fasting-induced torpor. **(A)** Time course study of torpor-activated neurons. Mice were fasted at 16°C from the onset of dark phase. The brains were collected at indicated time and the coronal sections were stained with anti-phsopho-S6 antibody. Top, a representative tracing of Tb in torpid mice. Bottom, immunolabeling of phospho-S6 in the DMH. Scale bar, 100μm. **(B)** Unilateral injection of the AAV-DIO eYFP into the DMH of Vgat-cre mice (left). Representative immunofluorescence images of the DMH from torpid Vgat-cre mice injected with AAV-DIO eYFP (right). Scale bar, 100μm. **(C)** The percentage of eYFP-positive or -negative DMH neurons in pshospho-S6-positive neurons in Vgat-cre mice injected with AAV-DIO eYFP. The number of cells in 2 or 3 of coronal brain sections from three torpid mice were counted. **(D)** Biilateral injection of AAV-DIO Kir2.1-zsGreen (top). Representative immunofluorescence image of the eGFP expression (bottom). Scale bar, 100μm. **(E)** Representative physiological tracings of Vgat-cre mice bilaterally injected with AAV-DIO eYFP or AAV-DIO Kir2.1 into the DMH upon fasting at cold. **(F)** Quantification of the change in Tb in mice injected with AAV-DIO eYFP(n=8) or AAV-DIO Kir2.1(n=7). ⊿Tb was defined as the mean Tb for the first 120 min minus the minimum Tb.^***^*P* <0.001 by two-tailed Student *t* -test between eYFP and Kir2.1. **(G)** Quantification of the change in VCO_2_ in mice injected with AAV-DIO eYFP (n=4) or AAV-DIO Kir2.1-zsGreen (n=4). ⊿VCO_2_ was defined as the percentage reduction in VCO_2_ by comparing the mean VCO_2_ for the first 120min to the minimum VCO_2_. ^*^*P* <0.05 by two-tailed Student *t* -test between eYFP and Kir2.1. **(H)** Bilateral injection of the AAV-DIO eYFP or Kir2.1 into the MPOA of Trpm2-cre mice (top). Representative immunofluorescence images of the expression of eYFP in the MPOA (bottom). Scale bar, 100μm. **(I)** Representative physiological tracings of Trpm2-cre mice bilaterally injected with AAV-DIO eYFP or AAV-DIO Kir2.1-zsGreen into the MPOA upon fasting at cold. **(J)** Quantification of the change in Tb in mice injected with AAV-DIO eYFP(n=8) or AAV-DIO Kir2.1-zsGreen(n=8). ⊿Tb was defined as the mean Tb for the first 120 min minus the minimum Tb. ^***^*P* <0.001 by two-tailed Student *t* - test between eYFP and Kir2.1. **(K)** Quantification of the change in VCO_2_ in mice injected with AAV-DIO eYFP (n=4) or AAV-DIO Kir2.1-zsGreen (n=4). ⊿VCO_2_ was defined as the percentage reduction in VCO_2_ by comparing the mean VCO2 for the first 120min to the minimum VCO2. ^*^*P* <0.05 by two-tailed Student *t* -test between eYFP and Kir2.1. **(L)** Unilateral injection of AAV-DIO eYFP or AAV-DIO ChR2-eYFP (top). Representative immunofluorescence image of the eYFP expression (bottom). Scale bar, 100μm. **(M)** Representative physiological tracings of Trpm2-cre mice bilaterally injected with AAV-DIO eYFP or AAV-DIO Kir2.1-zsGreen into the MPOA upon fasting at cold. Time course change in Tb during optogenetic stimulation (10Hz, 60min) of MPOA Trpm2-expressing neurons. n=4 per group. **(N)** Quantification of the change in Tb in mice injected with AAV-DIO eYFP (n=5) or AAV-DIO Kir2.1-zsGreen (n=5) upon optogenetic stimulation (488nm, 10Hz, 60min). ⊿Tb was defined as the mean Tb during 60min of pre-stimulation period minus the minimum Tb during optogenetic stimulation period. ^**^*P* <0.001 by two-tailed Student *t* -test between eYFP and ChR2. **(O)** The percentage of the change in locomotor activities in mice bilaterally injected with AAV-DIO eYFP (n=5) or AAV-DIO Kir2.1-zsGreen(n=5) upon optogenetic stimulation (10Hz, 60min). Total counts of locomotor activity for stimulation period was divided by total counts of pre-locomotor activity for stimulation period.^*^*P* <0.05 by two-tailed Student *t* -test between eYFP and ChR2.

We next injected AAV expressing Kir 2.1, an inwardly rectifying potassium ion channel that hyperpolarizes neurons^14^ in a Cre-dependent manner, bilaterally into the DMH of Vgat mice (Fig. 2D) to suppress neuronal activity. As a control, we injected AAV carrying cre-dependent eYFP bilaterally into the DMH of Vgat mice. Fasting at cold ambient temperature induced torpor in eYFP-expressing mice, with a maximum decrease in Tb of 9.7 ± 1.0 ° C (Fig. 2E and 2F). On the other hand, Kir 2.1 expressing mice did not enter torpor during fasting at cold ambient temperature (Fig. 2E and 2F). Instead, Tb decreased by 3°C in the first 2 hours and reached a plateau in the remaining recordings (Fig. 2E). The VCO_2_ in mice injected with Kir 2.1 was reduced to 58.1 ± 1.7%, whereas in control mice was reduced to 72.5 ± 5.0% (Fig. 2E and 2G). This could not be explained by differences in consumption and body weight prior to the experiment which were comparable between the eYFP and Kir 2.1-expressing groups(fig. S4). Next, we examined whether activation of DMH GABAergic neurons is sufficient to induce torpor. To this end, we injected Vgat-cre mice bilaterally with AAV expressing Cre-dependent Gq-coupled human M3 muscarinic receptor (hM3Dq)^15^, or eYFP as control, in the DMH. Three weeks after viral injection, DMH GABAergic neurons were chemically activated, but Tb did not decrease in any of the two groups (fig. S5). These results indicate that activation of DMH GABAergic neurons is necessary but not sufficient to induce torpor.

### Median Preoptic Trpm2 neurons receive projections from DMH and are necessary for torpor

DMH GABAergic neurons project extensively to the medial preoptic area, a critical brain region for thermoregulation^12,16,17^, and our brain-wide screen also showed that the preoptic area was also activated during torpor (Fig. 1E and 1F). In addition, Trpm2-expressing neurons in the MPOA detect hypothalamic temperature, and its activation decreases Tb^18^. These results led us to investigate the role of Trpm2-expressing neurons in the regulation of torpor. AAV carrying cre-dependent Kir 2.1 or eYFP was injected bilaterally into the MPOA of Trpm2-cre mice (Fig. 2H). Trpm2-cre mice expressing Kir 2.1 in the MPOA did not decrease Tb and VCO_2_ when fasted at 16 ° C in comparison to eYFP-expressing control mice(Fig. 2I, 2J, 2K), indicating that the activity of Trpm2-expressing neurons in the MPOA is necessary for initiating torpor. Food consumption and body weight did not differ between the eYFP and Kir 2.1-expressing groups one day before induction of torpor (fig. S6A and B). To investigate whether activation of Trpm2-expressing neurons is sufficient to initiate torpor, we injected unilaterally AAV carrying Cre-dependent channelrhodopsin 2 (ChR2) or eYFP and implanted an optic cannula into the MPOA of Trpm2-cre mice (Fig. 2L) and stimulated the cells using blue light. In mice injected with AAV-DIO ChR2, optogenetic stimulation decreased Tb by 7 ° C, whereas eYFP control mice were not affected by stimulation (Fig. 2M and 2N). Mice injected with ChR2 rapidly regained Tb when stimulation ceased (Fig. 2M). The locomotor activity of mice increased during optogenetic stimulation (Fig. 2M and 2O), in contrast to fasting-induced torpid mice, which stopped moving and exhibited a rounded posture (fig. S1). Taken together, these results indicate that the activity of Trpm2-expressing neurons in MPOA is essential and sufficient to induce torpor with respect to body temperature.

To elucidate the circuit controlling fasting-induced torpor, we performed anterograde and retrograde virus tracing of DMH GABAergic neurons and MPOA Trpm2-expressing neurons. We injected AAV carrying cre-dependent synaptophysin fused with mRuby^19^ unilaterally into the DMH of Vgat-cre (Fig. 3A). Labeling in the medial preoptic area, periaqueductal gray (PAG), and raphe pallidus (RPa) indicated synaptic innervation of DMH GABAergic neurons onto these areas (Fig. 3B). We then injected unilaterally AAV carrying Cre-dependent rabies G and TVA, followed by injection of EnvA pseudotype eGFP-expressing delta-G rabies virus into the DMH of Vgat mice, to perform retrograde virus tracing (Fig. 3C and 3D). We found labeling in the BNST, median preoptic area (MnPO), medial preoptic area, PVN, SCN, PVp, TMN, and RPa formed pre-synapses onto GABAergic neurons in the DMH (Fig. 3E). Next, we examined the input/output structure of Trpm2-expressing neurons in MPOA. Unilateral injection of AAV carrying Cre-dependent synaptophysin-mRuby (Fig. 3F) into the MPOA of Trpm2-cre mice revealed its projections to the periventricular nucleus, DMH, and TMN (Fig. 3G). Cre-Dependent retrograde rabies tracing (Fig. 3H and 3I) also revealed that BNST, median preoptic area, PVT, PVN, Arc, medial amygdala (MeA), PVp, and RPa formed pre-synapses onto Trpm2-positive neurons in MPOA (Fig. 3J). BNST, PVN, PVp and RPa projected to both DMH and MPOA. Notably, the BNST and PVN were activated in torpid mice but not in control groups (Fig. 1E). The reciprocal connections between the MPOA and DMH suggest cooperative regulation of torpor by these regions. Fig. 3K illustrates a postulated schematic diagram of the neural circuitry that regulates fasting torpor.

**Fig. 3.**
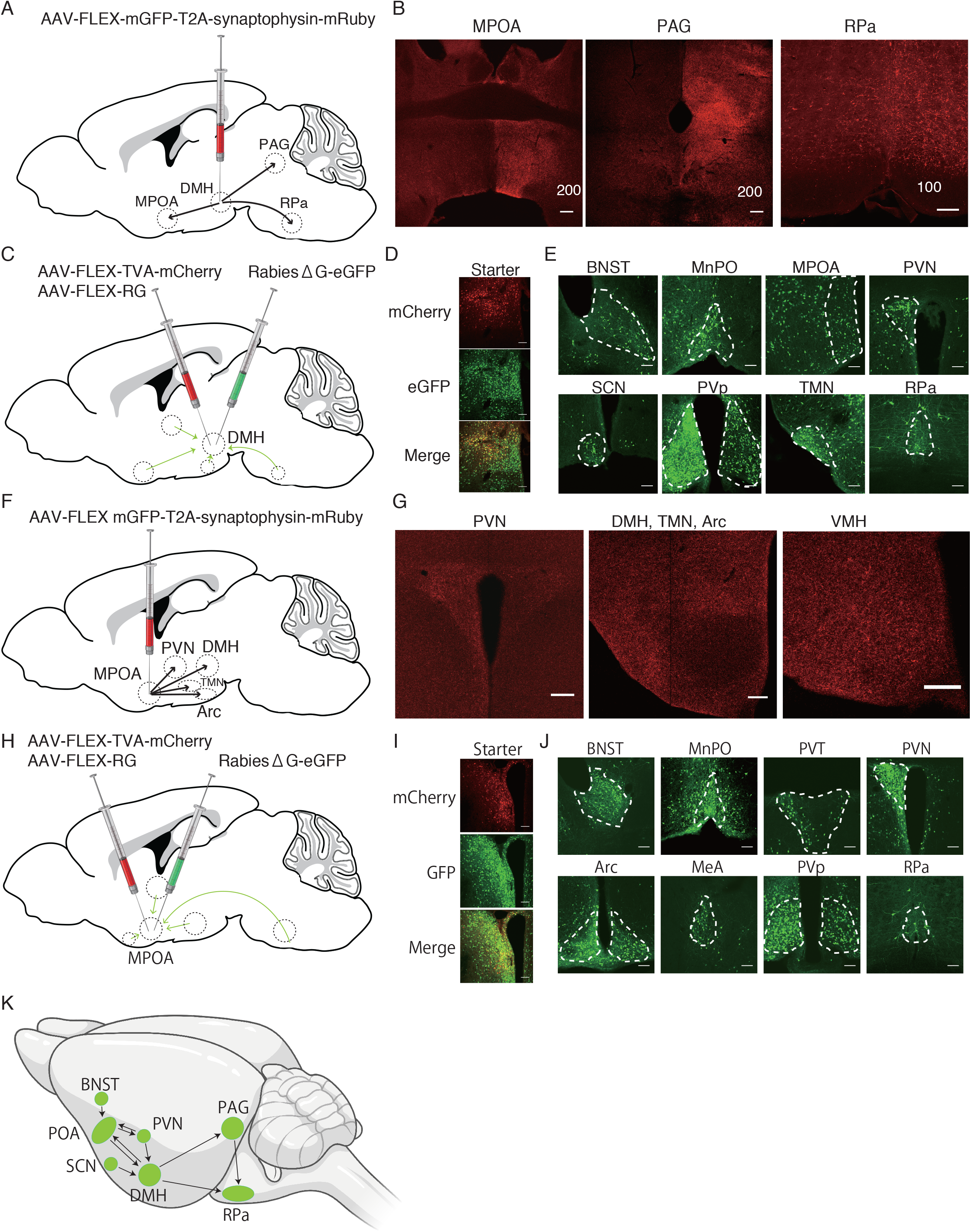
Viral tracing of hypothalamic torpor-activated neurons. **(A)** AAV-DIO synaptophysin-mRuby was unilaterally injected into the DMH of Vgat-cre mice. **(B)** mRuby-positive puncta from DMH-Vgat neurons in the medial preoptic area, periaqueductal grey and raphe pallidus. Scale bar, 100 (right) or 200μm (left and middle). **(C)** Experimental design of the trans-synaptic retrograde tracing. Three weeks after the injection of AAV-FLEX TVA-mCherry and AAV-FLEX RG, EnvA pseudotyped RG-deleted rabies virus expressing eGFP was injected into the DMH of Vgat-cre mice. **(D)** Representative images of mCherry and eGFP-double positive starter cells in the DMH. Scale bar, 100μm. **(E)** Representative images of upstream eGFP-positive cells providing direct inputs onto DMH Vgat neurons, replicated independently with similar results in three mice. Scale bar, 100μm. **(F)** AAV-DIO synaptophysin-mRuby was unilaterally injected into the MPOA of Trpm2-cre mice. **(G)** mRuby-positive puncta from MPOA-Trpm2 neurons in the PVN, DMH, Arc and VMH. Scale bar, 100 (right) or 200μm (left and middle). **(H)** Experimental design of the trans-synaptic retrograde tracing. Three weeks after the injection of AAV-FLEX TVA-mCherry and AAV-FLEX RG, EnvA-pseudotyped RG-deleted rabies virus expressing eGFP was injected into the MPOA of Trpm2-cre mice. **(I)** Representative images of mCherry and eGFP-double positive starter cells in the MPOA. Scale bar, 100μm. **(J)** Representative images of upstream eGFP-positive cells providing direct inputs onto MPOA Trpm2 neurons, replicated independently with similar results in three mice. Scale bar, 100μm. **(K)** Schematic diagram of the neural circuitry that regulates fasting-induced torpor.

## Discussion

While torpor is an important protective behavior against harsh environments, active suppression of metabolic demand during torpor may improve the treatment of ischemic injury^20,21^. Thus, investigating the neural mechanisms underlying torpor may confer therapeutic benefits beyond ecological contexts. In this study, we mapped neurons activated during fasting-induced torpor by whole-brain immunolabeling with an active neuron marker. As a proof-of-concept, we determined the essential roles of DMH GABAergic neurons and MPOA Trpm2-expressing neurons in the initiation of torpor. Fasting-induced torpor is regulated by circadian rhythms because it begins in the late dark phase, regardless of when fasting begins^22^. However, it is not yet known which neuronal populations integrate internal and external signals such as fasting, ambient temperature, and circadian rhythm signals to regulate torpor. DMH GABAergic neurons have active inputs from the arcuate nucleus, a brain region that senses fasting signals from peripheral tissues^23^, as well as the suprachiasmatic nucleus, the center of circadian rhythms^24^. Thus, this region may be involved in controlling the onset timing of torpor. Also, in fasting-induced torpor, decrease and re-increase of body temperature follow the ultradian cycle. Enoki *et al*. showed that the PVN, a common upstream region of DMH GABAergic and MPOA Trpm2-expressing neurons, had a firing pattern that followed ultradian rhythm^25^. It will be interesting to examine whether the activity of neurons in the PVN is involved in the timing of torpor and wakefulness.

Recently, it was shown that activation of preoptic neurons expressing pyroglutamylated RFamide peptide (Qrfp)^26^or adenylate cyclase activating polypeptide 1 (Adcyap1)^27^ induced torpor-like decrease in Tb. Suppression of neurotransmission from these neurons disrupted the fast decrease in body temperature during torpor. Interestingly, the decrease in body temperature observed was gradual, suggesting that there is a mechanism that regulates hypothermia during fasting-induced torpor independent of the activity of these neurons. Furthermore, preoptic neurons expressing the estrogen receptor were reported to be involved in the maintenance, but not the initiation, of fasting-induced torpor^28^. In contrast, our study demonstrates that MPOA Trpm2-expressing neurons are necessary for triggering torpor. Therefore, these data indicate that body temperature and metabolism during torpor are cooperatively regulated by multiple populations of the preoptic area.

Torpor is a complex behavior to save energy consumption, and animals during torpor not only reduce heat production and body temperature, but also reduce heat loss from the body surface by curling up and vasoconstriction. Optogenetic activation of Trpm2-positive cells did indeed lower body temperature but did not reproduce the unique curled posture during torpor. These results suggest that the full reproduction of fasting-induced torpor requires the activity of other circuits. The whole-brain screen used in this study found neurons that were activated during torpor in addition to the hypothalamus, and these neurons may control other torpor characteristics, such as the torpor posture.

Taken together, our results highlight the utility of unbiased whole-brain screening in interrogation of the neural circuits underlying innate behavior and provide the groundwork for future studies of fasting-induced torpor, with potential translational implications metabolic trauma medicine and space travel.

## Materials and Methods

### Animals

Trpm2^tm1.1(icre)Jsmn^/J(TRPM2-Cre; JAX #029047), Vglut2 or Slc32a1^tm2(cre)Lowl^/J (Vgat-Cre; JAX #016962) were obtained from Jax. Female mice aged 10 ∼ 16 weeks were used at the start of the experimental procedures. Mice were housed in a Plexiglas recording chambers with a reversed circadian cycle (12 hours dark-light cycle). Food and water were available ad libitum. All experiments were performed in accordance with the guidelines described in the US National Institutes of Health Guide for the care and Use of Laboratory Animals and approved by Stanford University’s Administrative Panel on Laboratory Animal care and the Institutional Animal Care and Use Committees of the Research Institute of Environmental Medicine, Nagoya University.

### Viral injection

Animals were anesthetized by intraperitoneal injection of ketamine (100 mg/kg) and xylazine (10 mg/kg). Buprenorphine (0.5 mg/kg) was administered subcutaneously to the animals as post-operational analgesia. Recombinant AAV was injected either unilaterally or bilaterally into the medial preoptic hypothalamus of Trpm2-cre mice (AP = 0 mm; ML = ± 0.3 mm; DV = -5.3 mm) or the dorsomedial hypothalamus of Vgat-cre or Vglut2 mice (AP = -1.75 mm; ML = ± 0.32 mm; DV = -5.20 mm). The injections were performed at a rate of 50 nl/min (Total 400 nl) via a 33 gauge needle (Hamilton) attached to a 5.0 ml Hamilton syringe. The titers of the viruses used were as follows; AAV-DJ EF1α DIO ChR2-eYFP (4.0 × 10^12^ gc/ml), AAV-DJ CMV DIO Kir 2.1-zaGreen (1.0 × 10^13^ gc/ml), AAV-DJ EF1α DIO eYFP (6.0 × 10^12^ gc/ml), AAV-DJ EF1α DIO synaptophysin-mRuby -2 A-GFP (2.6 × 10^13^ gc/ml), AAV5 EF1α DIO hM3Dq mCherry ((3.4×10^12^ gc/ml). Two weeks after viral injection, mice for optogenetic experiments were subjected to unilateral surgical implantation of a fiber optic cannula (200 μm; Doric Lenses, Inc.) above the preoptic hypothalamus (AP = 0 mm; ML=-0.3 mm; DV=-5.25mm). For retrograde rabies tracking, a mixture of AAV-DJ EF1α FLEX TVA-mCherry (4.9×10^12^ gc/ml)) and AAV-DJ CAG FLEX RG (1.3×10^12^ gc/ml) was injected into the medial preoptic hypothalamus of TRPM2-cre mice or the dorsomedial hypothalamus of Vgat-cre mice, respectively. Three weeks after the first injection, EnvA pseudotype RG-deleted rabies virus expressing eGFP(1.0×10^8^ TU/ml) was injected into the area where starter AAV was injected. The brains were collected from the animals one week after rabies virus injection.

### Torpor induction

Mice were intraperitoneally implanted with a temperature and activity telemetry transmitter (DSI; TA-F 10). After a recovery period of at least 7 days from surgery, the mice were transferred from the home cage to a temperature-controlled cage (16 ° C or 31 ° C) at the onset of the dark period of the reversed circadian cycle (12 hour dark-light cycle). The mice were then fasted or ad libitum fed for 24 hours. Data of core body temperature and locomotor activity were collected and analyzed by Dataquest ART (DSI).

### Measurement of CO2 production

Animals were housed in clear plexiglass Oxymax^®^-CLAMS chambers. Each chamber was equipped with an analog infrared CO2 sensor for analog measurement of CO2 concentrations using custom data acquisition software (DFRobot.com, SKU SEN0219). CO2 accumulation relative to ambient concentration were measured once every 5 minutes prior to chamber perfusion with room air and normalized to animal body mass.

### Immunolabeling of whole brain and iDISCO brain clearing

Mice were anesthetized with ketamine/xylazine and trans-cardinally perfused with 1×PBS (pH 7.4), followed by 4% paraformaldehyde in PBS. Fixed samples were washed with PBS for 3 × 30min, then dehydrated in 20%, 40%, 60%, 80%, 100% methanol (in water) for 1 hr each. Samples were further washed with 100% methanol for 1 hr and then chilled at 4°C. Samples were then incubated in 66% dichloromethane/33% methanol for overnight at room temperature, then washed with 100% methanol for 10 min twice, then chilled at 4°C. Samples were then bleached with 5% H_2_O_2_ in methanol (1vol 30% H2O2/5vol methanol) for overnight at 4°C. After bleaching, samples were rehydrated in 80%, 60%, 40%, 20% methanol (in water) for 1 hr each and then washed with PBS for 1 hr at room temperature. Samples were then washed with PBS/0.2% Triton X-100 (PTx.2) and permeabilized in PBS/0.2% Triton X-100/20% DMSO/0.3M glycine for 48 hr at 37°C. Samples were then blocked in PBS/0.2% Triton X-100/10% DMSO/6% Donkey serum at 37°C for 48 hr and incubated in primary antibody dilutions (rabbit anti-phospho-S6, 1:300) in PBS/0.2% Tween-20 containing 10μg/ml heparin (PTwH) added with 5% DMSO and 3% Donkey serum for 14 days at 37°C. Samples were then washed in PTwH for 1day and incubated with secondary antibody (Alexa Fluor 647 donkey anti-goat IgG, Invitrogen, 1:500) in PTwH/3% Donkey serum for 14 days at 37°C. Samples were then washed in PTwH for 1 day. Immunolabeled samples were then dehydrated in 20%, 40%, 60%, 80%, 100% methanol (in water) for 1 hr each and washed in 66% dichloromethane/33% methanol for 3 hr and with 100% dichloromethane for 15 min twice at room temperature. Samples were then cleared in dibenzyl ether.

### Light-Sheet Imaging

Optically cleared brain samples were imaged in the sagittal direction (right lateral side up) on a light sheet microscope (Ultramicroscope II, LaVision Biotec) equipped with a sCMOS camera and a 2 ×/0.5 objective lens. Samples were scanned at a 3 μm step size using a continuous-light sheet scanning method with the included contrast mixing algorithm for 640 nm and 595 nm channels.

### ClearMap Analysis

Immunoreactive cells in the optically cleared brain samples were mapped and quantified using ClearMap software^11^. Cell detection is tailored to the cell body and uses background subtraction via morphological opening, followed by a series of filters, morphological manipulation, and 3D peak detection. 700 voxels was used as the threshold for cell size. The background was removed by subtraction of the morphological opened image by a disk-shaped structural element with a main axis of 7 pixels in diameter. The heatmaps were created by summing spheres of uniform intensity, 375 μm in diameter, centered on each cell, and the p-value density maps (Fig. 1F) were created from the heatmaps. Samples were mapped using the average autofluorescence STPR brain (Kim et al., 2015) registered to the Allen Brain Institute 25 (μm/pixel) map and companion annotation map (http://alleninstitute.org/).

### Immunohistochemistry

Mice were anesthetized with ketamine/xylazine and trans-cardinally perfused with 1×PBS, pH 7.4, followed by 4% paraformaldehyde in PBS. Brains were collected, post-fixed for overnight at 4°C, and then cryoprotected in 30% sucrose dissolved in PBS containing 0.1% sodium azide for more than 48 hr at 4°C. Brain sectioning was performed on a cryostat (Leica Microsystems) at a thickness of 60 μm. Sections were rinsed in PBS with 0.3% triton X-100 (PBST) and blocked with 2% bovine serum albumin in PBS for 1hr at room temperature. Sections were then incubated in blocking buffer containing primary antibodies overnight at room temperature. After 3 × 5min washes in PBST, sections were incubated in blocking buffer containing secondary antibodies for 6 hr at room temperature. After the incubation, sections were washed in PBST three times and then mounted onto MAS-GP microscope slides (Matsunami) and coverslipped with Fluoroshield Mounting Media (Abcam). Images were collected on a confocal microscope (Zeiss LSM 710, Zen Software). Quantification of colocalization was performed on serial sections from approximate bregma -5.12 to -5.76 from three mice. The antibodies used were as follows; rabbit anti-phospho-S6 244/247 (1:500, ThermoFisher #44-923G), goat anti-cFos (1:500, Santa Cruz #sc-52G), Alexa Fluor 594 donkey anti-rabbit IgG (1:1000, Invitrogen #A21207), Alexa Fluor 647 donkey anti-goat IgG (1:1000, Invitrogen #A21447).

### Statistics

Sample sizes were chosen based upon Mead’s rule and previous publications of the study of thermoregulation. Data distribution was assumed to be normal, but not formally tested. We analyzed all data using Prism 6.0 (Graphpad Software) and data are presented as mean±s.e.m. We used Prism 6.0 and Adobe Illustrator CS6 (Adobe Systems) to prepare the figures.

## Acknowledgements

We thank all members of L.d.L’s Lab for critical feedback in the preparation of this manuscript. H.Y. was supported by Uehara memorial foundation research fellowship and the Lotte Shigemitsu Prize. K.M. was supported by the National Heart Lung and Blood Institute grant 5T32HL110952-07. L.d.L. is supported by National Institutes of Health Grants AG047671, MH087592, MH102638. The light sheet Ultramicroscope II is supplied by NIH shared equipment grant 1S10OD025091-01.

---

**Supplemental Fig. 1**. Locomotor activity and characteristic posture during torpor. (A) Representative tracings of body temperatures (Tb) and locomotor activity in mice at 16°C with fasted for 24-hr recording. (B) Image of a torpid mouse.

**Supplemental Fig. 2**. The onset timing of torpor at 16°C with fasted. The time when mice fasted at 16°C enter torpor were plotted (n=10, mean=437.4min).

**Supplemental Fig. 3**. Automated analysis of pS6+ cell distribution in mouse brains collected under different conditions at ZT20 (n=4). The number of pS6 + cells was counted per each voxel. Panels show the reference annotation, the averaged cell density maps for each group, and p-value maps for ventral lateral septal nucleus (LSV), bed nucleus of stria terminalis (BNST) and anterior commissure, anterior(aca). The cell density from each group were compared with that of control mice fed at thermoneutrality by the t-test and their p-values (Magenta: fasted at 31°C, Red: fed at 16°C、Green: fasted at 16°C) were mapped. Scale bar, 300μm.

**Supplemental Fig. 4**. Body weight and food intake of Vgat-cre mice injected with AAV-DIO eYFP (n=4) or AAV-DIO Kir2.1-zsGreen (n=4). n.s. *P*>0.05 by two-tailed Student *t* -test between eYFP and Kir2.1.

**Supplemental Fig. 5**. Time course change in Tb during chemogenetic activation of DMH Vgat-expressing neurons. Vgat-cre mice were injected with AAV-DIO mCherry (n=4) or AAV-DIO hM3Dq mCherry (n=4) into the DMH. At 60min, 1mg/kg of CNO or saline was intraperitoneally injected into mice.

**Supplemental Fig. 6**. Body weight and food intake of Trpm2-cre mice injected with AAV-DIO eYFP (n=4) or AAV-DIO Kir2.1-zsGreen (n=4). n.s. *P* >0.05 by two-tailed Student *t* -test between eYFP and Kir2.1.

**Supplemental Movie. 1**. Automated analysis of pS6+ cell distribution in mouse brains collected under different conditions at ZT20 (n=4). The number of pS6 + cells was counted per each voxel. The cell density from each group were compared with that of control mice fed at thermoneutrality by the t-test and their p-values (Magenta: fasted at 31°C, Red: fed at 16°C、Green: fasted at 16°C) were mapped.

## References

1. Carey, H. V., Andrews, M.T. & Martin, S.L. Physiol. Rev. 83, 1153–1181 (2003).

2. Ruf, T. & Geiser, F. Biol. Rev. 90, 891–926 (2015).

3. Hudson, J.W. & Scott, I.M. Physiol. Zool. 52, 205–218 (2016).

4. Heller, H.C. & Hammel, H.T. Comp. Biochem. Physiol. -- Part A Physiol. 41, 349–359 (1972).

5. Tamura, Y., Shintani, M., Nakamura, A., Monden, M. & Shiomi, H. Brain Res. 1045, 88–96 (2005).

6. Jinka, T.R., Toien, O. & Drew, K.L. J. Neurosci. 31, 10752–10758 (2011).

7. Iliff, B.W. & Swoap, S.J. Am. J. Physiol. Integr. Comp. Physiol. 303, R477–R484 (2012).

8. Knight, Z.A. et al. Cell 151, 1126–1137 (2012).

9. Renier, N. et al. Cell 159, 896–910 (2014).

10. Goldowitz, D. Genes, Brain Behav. 9, 128–128 (2010).

11. Renier, N. et al. Cell 165, 1789–1802 (2016).

12. Nakamura, K. & Morrison, S.F. J. Physiol. 586, 2611–2620 (2008).

13. Geerling, J.C. et al. Am. J. Physiol. - Regul. Integr. Comp. Physiol. (2016).doi:10.1152/ajpregu.00094.2015

14. Hibino, H. et al. Physiol. Rev. 90, 291–366 (2010).

15. Roth, B.L. Neuron 89, 683–694 (2017).

16. Tan, C.L. et al. Cell 167, 47–59.e15 (2016).

17. Yu, S. et al. J. Neurosci. 36, 5034–5046 (2016).

18. Song, K. et al. Science (80-.). 353, 1393–1398 (2016).

19. Beier, K.T. et al. Cell 162, 622–634 (2015).

20. Dave, K.R., Christian, S.L., Perez-Pinzon, M.A. & Drew, K.L. Comp. Biochem. Physiol. - B Biochem. Mol. Biol. 162, 1–9 (2012).

21. Cerri, M. Annu. Rev. Physiol. 79, 167–186 (2016).

22. Van Der Vinne, V., Bingaman, M.J., Weaver, D.R. & Swoap, S.J. J. Exp. Biol. (2018).doi:10.1242/jeb.179812

23. Andermann, M.L. & Lowell, B.B. Neuron (2017).doi:10.1016/j.neuron.2017.06.014

24. Hastings, M.H., Maywood, E.S. & Brancaccio, M. Nat. Rev. Neurosci. (2018).doi:10.1038/s41583-018-0026-z

25. Wu, Y.E. et al. Proc. Natl. Acad. Sci. U. S. A. (2018).doi:10.1073/pnas.1804300115

26. Takahashi, T.M. et al. Nature (2020).

27. Hrvatin, S. et al. Nature (2020).doi:10.1038/s41586-020-2387-5

28. Zhang, Z. et al. Nat. Commun. (2020).doi:10.1038/s41467-020-20050-1

