## Supplementary figures and images for "Brain-wide mapping of neuronal architecture controlling torpor"

### Supplemental Figure.1

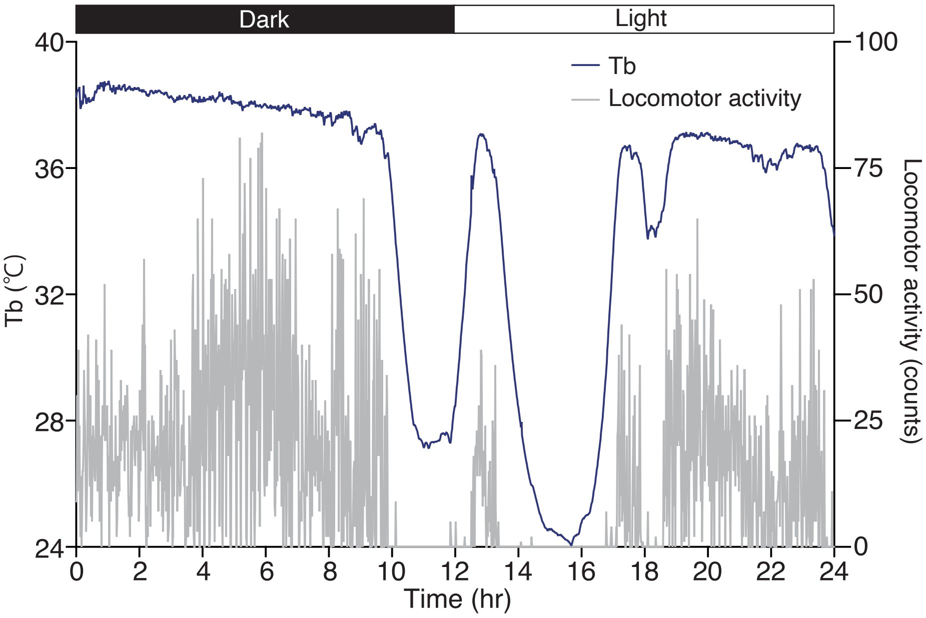

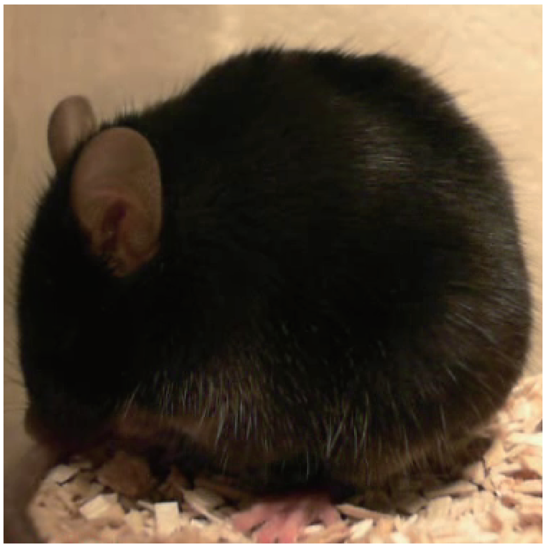

### Supplemental Figure.2

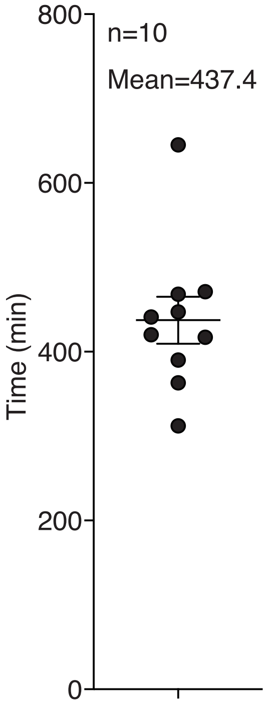

### Supplemental Figure.3

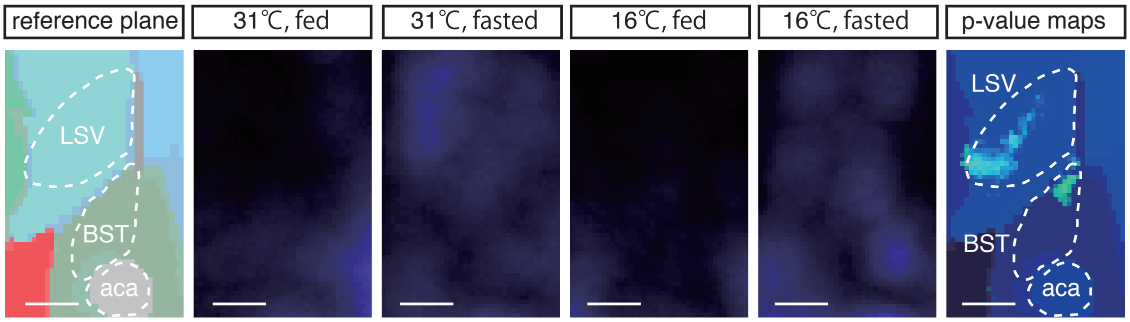

### Supplemental Figure.4

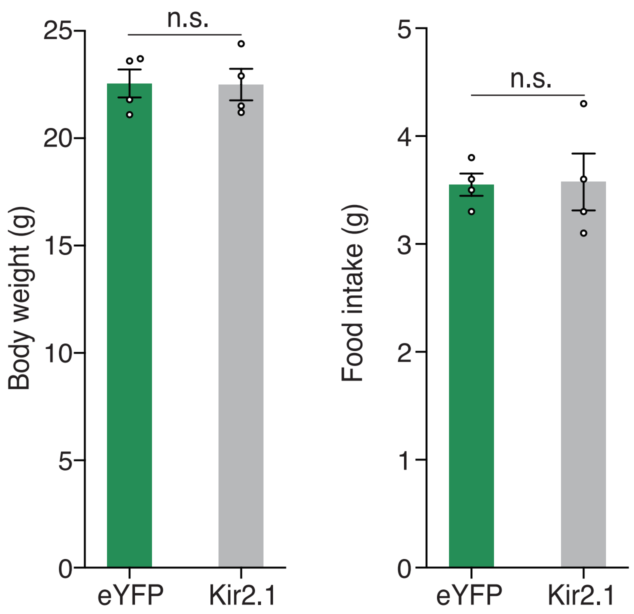

### Supplemental Figure.5

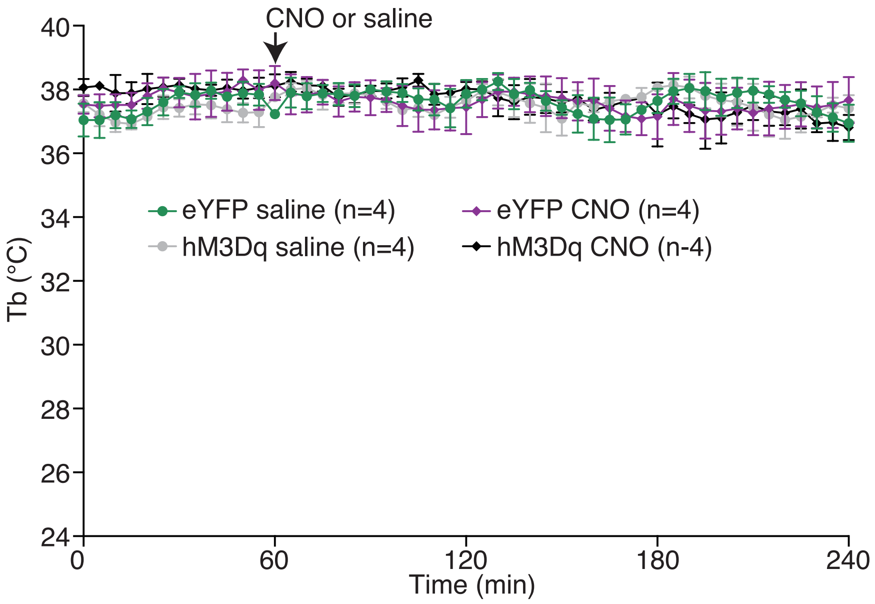

### Supplemental Figure.6

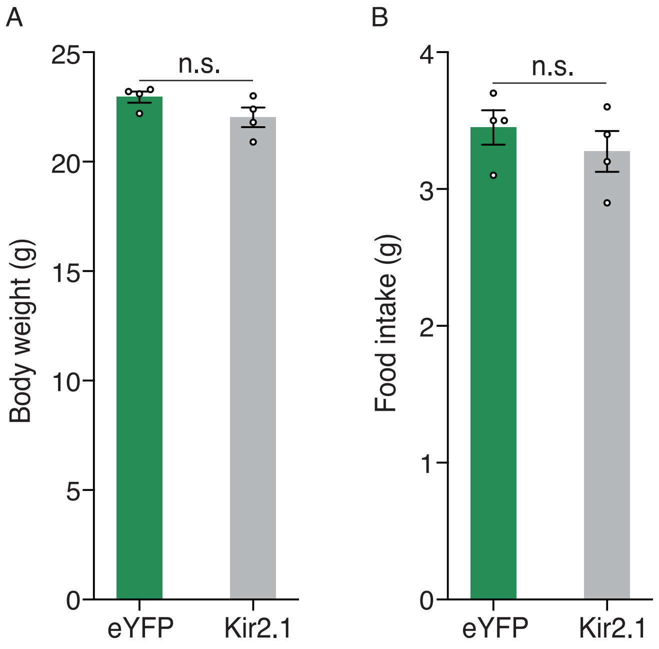
